# Multilevel clinical fingerprinting: uncovering longitudinal changes in the functional connectome of the brain along the migraine cycle

**DOI:** 10.1101/2024.05.02.592281

**Authors:** Inês Esteves, Ana R. Fouto, Amparo Ruiz-Tagle, Gina Caetano, Rita G. Nunes, Nuno A. da Silva, Pedro Vilela, Raquel Gil-Gouveia, Isabel Pavão Martins, César Caballero-Gaudes, Patrícia Figueiredo

## Abstract

Migraine is a common neurological disorder characterized by recurrent headache episodes alternating with symptom-free periods, which has been associated with alterations across large-scale functional brain networks albeit with variable findings. Critically, despite the cyclic nature of the disorder, longitudinal studies spanning the various phases of the migraine cycle are scarce. Here, we leverage the identifiability of individual functional connectomes (FC) to investigate changes along the migraine cycle. For this purpose, we employ a case-control longitudinal design to study a group of 10 patients with episodic menstrual or menstrual-related migraine without aura, in the 4 phases of their spontaneous migraine cycle (preictal, ictal, postictal, interictal), and a group of 14 healthy controls in corresponding phases of the menstrual cycle, using resting-state fMRI. We propose a novel multilevel clinical fingerprinting approach to analyse the differential FC identifiability within-subject, as well as within-session and within-group. The individual FC matrices are then reconstructed with 19 principal components maximizing identifiability at all levels, and analyzed with Network-Based Statistic to identify significant changes in FC strength. We observe decreased FC identifiability for patients in the preictal phase relative to controls, which increases with the progression of the attack and becomes comparable to controls in the interictal phase. Regarding the FC strength, is increased in the ictal and postictal phases relative to controls across several networks. Our novel multilevel clinical fingerprinting approach captures FC variations along the migraine cycle in a case-control longitudinal study, bringing new insights into the cyclic nature of the disorder.

## 1. Introduction

Migraine is a cyclic condition characterized by recurrent headache attacks, accompanied by sensory and cognitive disruptions, alternated with headache-free periods. It is defined by four distinct phases: preictal (before the headache begins), ictal (the headache phase lasting 4–72 hours), postictal (following headache cessation) and interictal (in-between the attacks) (Peng & May, 2020). Migraine is considered the second cause of disability worldwide and the first cause for women within 15-49 years old (Steiner et al., 2020), and its prevalence is two to three times higher in women than in men. Furthermore, menstrual migraine and menstrual-related migraine are among the most common subtypes for female patients (20-25%)(Vetvik & MacGregor, 2021), with menstrual-related attacks being considered more debilitating (Pavlović et al., 2015). Over the past decades, magnetic resonance imaging (MRI) has provided insights beyond clinical assessment, revealing structural and functional brain alterations in migraine (Colombo et al., 2019; Messina et al., 2022, 2023).

Changes in functional connectivity across large-scale brain networks, referred to as functional connectome (FC), have been observed in migraine patients using resting-state functional magnetic resonance imaging (fMRI); however, findings are highly variable across studies (Chou et al., 2023; Schramm et al., 2023; Skorobogatykh et al., 2019). Notably, most reports pertain to the interictal phase and studies examining the migraine cycle are limited, with a remarkable scarcity of investigations during the peri-ictal phases (preictal, ictal and postictal). In fact, studying the phases surrounding the attack is more challenging due to practical hurdles, such as the unpredictability of the attack and patient discomfort (Tolner et al., 2019). In a comprehensive review spanning fMRI studies from 2014 to 2021, only a small fraction (9 out of 114) included data partially collected during the ictal phase, and often the cycle phases were poorly defined (Schramm et al., 2023). Moreover, only 14 FC studies sampled more than one phase, and of these only 9 were longitudinal (Amin et al., 2018; Araújo et al., 2023; Filippi et al., 2022; Marciszewski et al., 2018; Meylakh et al., 2018; Schulte, Menz, et al., 2020; Schulte & May, 2016; Stankewitz et al., 2021; Stankewitz & Schulz, 2022), with some of them relying on the same sample and including 2 case studies. Furthermore, only 3 of the longitudinal studies sampled all migraine cycle phases and they did not compare them with healthy controls. Results were highly variable, with studies reporting increases and decreases, as well as no changes in FC across different networks. Importantly, despite having populations with mostly or only female participants, only a few of these studies indicate how the menstrual cycle was controlled for although it has been shown to influence FC (Dubol et al., 2021).

In an impactful work, FC has demonstrated the ability to identify unique profiles of individual subjects (Finn et al., 2015). Functional connectomes exhibit stability over time, irrespective of the task a subject is engaged in, resembling a functional fingerprint and enabling subject identification (Finn et al., 2015). Dimensionality reduction approaches can be used to further improve subject identifiability, resulting in higher brain-behaviour relationships (Amico & Goñi, 2018; Svaldi et al., 2021), and opening the door to further investigate the link to cognitive and clinical variables in the context of diseases. Building on the framework proposed in Amico & Goñi (2018), Sorrentino et al. (2021) introduced the concept of clinical connectome fingerprinting, which has been applied to FC data collected using magnetoencephalography (MEG) in different pathologies (Cipriano, Troisi Lopez, et al., 2023; Romano et al., 2022; Sorrentino et al., 2021; Troisi Lopez et al., 2023). A slightly different approach was applied to the study of psychosis patients using fMRI (Tepper et al., 2023). In general, these studies found reduced identifiability in patients compared with healthy controls, and that the differences in relation to controls can predict individual clinical features. To our knowledge, no study has yet employed a connectome fingerprinting approach to study patients longitudinally over more than 2 sessions, while also differentiating them from controls.

Here, we leverage the identifiability of individual functional connectomes (FC) to investigate changes along the migraine cycle. For this purpose, we employ a case-control longitudinal design to study a group of patients with episodic menstrual or menstrual-related migraine without aura, in the four phases of their spontaneous migraine cycle (preictal, ictal, postictal, interictal), and a group of healthy controls in corresponding phases of the menstrual cycle. We develop a novel multilevel clinical fingerprinting approach to investigate whether the migraine cycle influences individual FC fingerprints, relative to menstrual controls.

## 2. Materials and Methods

### 2.1. Participants

Data was acquired from 18 female patients suffering from episodic migraine without aura (M group), diagnosed according to the criteria of the 3rd edition of the International Classification of Headache Disorders (ICHD-III) (International Headache Society, 2018), with ages comprised between 18 to 55 years old. In addition, participants in the cohort experienced menstrual-related migraine attacks. The patients were otherwise healthy, without any diagnosed condition significantly hindering active and productive life, and with a life expectancy exceeding 5 years. None of them was receiving treatment with psychoactive drugs, including anxiolytics, antidepressants, anti-epileptics, and any migraine prophylactics. We aimed to assess patients in 4 sessions: preictal (M-pre), ictal (M-ict) and postictal (M-post) phases (around menses); and interictal phase (M-inter) (post-ovulation). For the interictal session, participants were required to be free from pain for at least 48 hours, with confirmation of the absence of a migraine attack obtained 72 hours post-scan. Of the 18 patients, only 10 (age 34.9±8.9 years) completed the 4 sessions and those were the ones included in the analysis. Several clinical features were also collected for the M group: Migraine Onset (19.3±6.0 years), Disease Duration (16.6±10.1 years), Attack Frequency (2.6±2.0 per month), Attack Duration (41.4±21.8 hours) and Attack Pain Intensity (6.7±1.1, from a scale 1-10). Furthermore, we collected data from 16 healthy controls (HC group) matched for gender, age, contraceptive use, and menstrual phase at the time of scanning. One of them was excluded to incidental findings and another one due to technical issues during the recording, and therefore the final sample was composed of 14 participants (age 34.9±8.9 years), They were assessed in two phases of the menstrual cycle so as to match the peri-ictal and interictal phases of the patients, yielding respectively the perimenstrual phase (approximately 5 days before or after the menses) and the post-ovulation phase (approximately on day 19 of the menstrual cycle). The experimental protocol is schematically represented in Figure 1.

**Figure 1.**
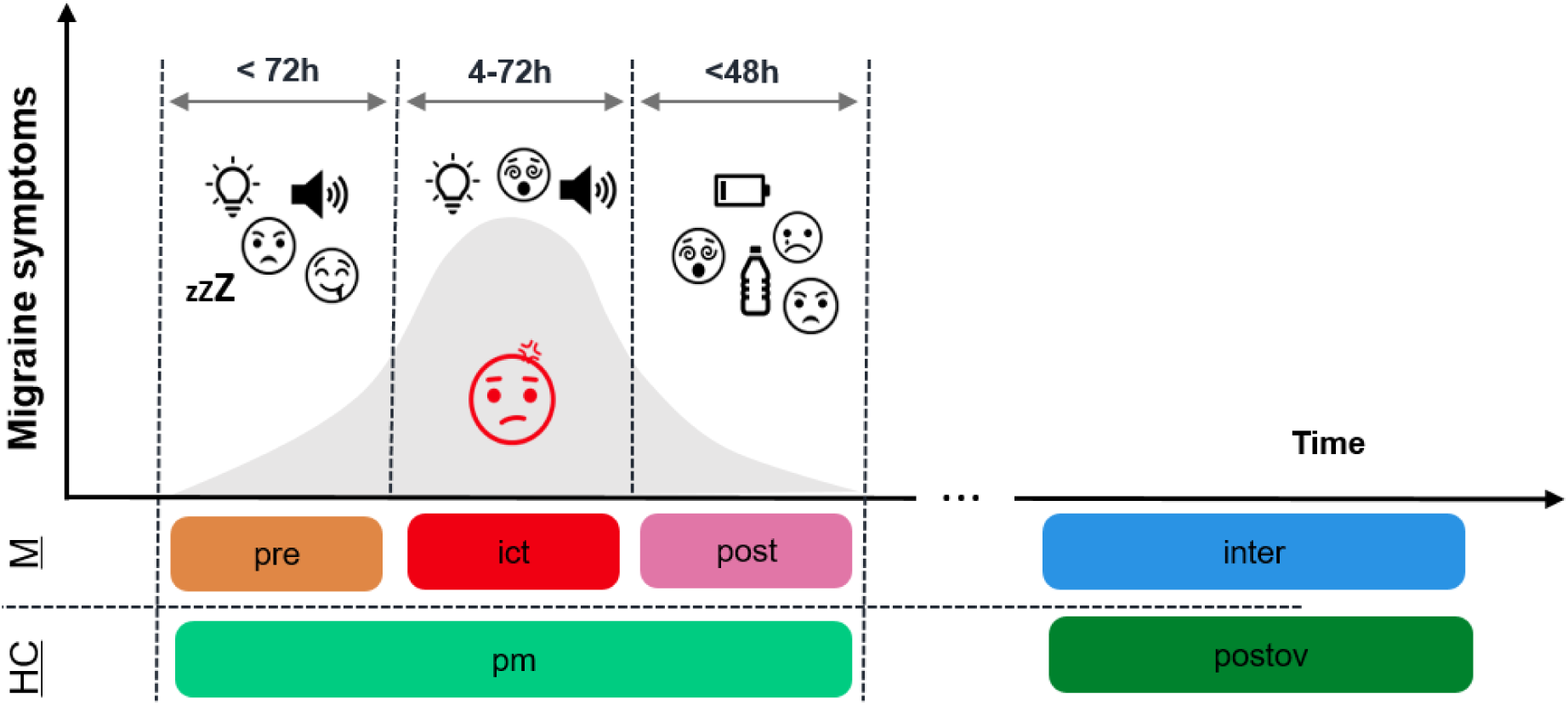
Definition of preictal (M-pre), ictal (M-ict), postictal (M-post) and interictal (M-inter) phases for the migraine patients (M) and the corresponding phases for the healthy controls (HC), matched for the menstrual cycle, perimenstrual (HC-pm) and post-ovulation (HC-postov). The symptoms illustrated across the migraine cycle are only present for the M group and include sleepiness, food cravings, increased sensitivity to light/sound, mood changes, cognitive dysfunction, fatigue, and thirst.

### 2.2. MRI data acquisition

The MRI data was acquired using a 3T Siemens Vida system, using a 64-channel RF coil. Functional images were obtained using a T2*-weighted gradient-echo Echo Planar Imaging (EPI) with the following parameters: TR/TE=1260/30ms, flip angle = 70°, in-plane generalized autocalibrating partially parallel acquisition (GRAPPA) acceleration factor = 2, simultaneous multi-slice (SMS) factor = 3, 60 slices, 2.2 mm isotropic voxel resolution. T1-weighted structural images were acquired with a magnetization-prepared rapid gradient echo (MPRAGE) sequence, with TR = 2300 ms, TE = 2.98 ms, inversion time (TI) = 900 ms, and 1 mm isotropic voxel resolution. Fieldmap magnitude and phase images were obtained using a double-echo gradient echo sequence (TR = 400.0 ms, TE = 4.92/7.38 ms, voxel size: 3.4 × 3.4 × 3.0 mm^3^, flip angle = 60°). Participants were instructed to keep their eyes open, looking at a black screen, and to not think in anything in particular, during 7 min (333 fMRI volumes). To mitigate acoustic noise exposure and minimize head motion, earplugs were used and small cushions were placed beneath and on the sides of the head of the subject.

### 2.3 MRI data preprocessing

The MRI data was preprocessed using FSL’s tools (Jenkinson et al., 2012).

Structural images were preprocessed, including nonbrain tissue removal using Brain Extraction Tool (BET) and bias field correction using FAST. Each strcutrual image was registered to the subject’s functional space (reference volume) as well as the standard MNI space, using FLIRT and FNIRT. White matter (WM) and cerebrospinal fluid (CSF) masks were derived by tissue segmentation using FAST, and subsequently registered to the functional space. For the CSF mask, an additional intersection with the ventricles (originally in the MNI space and transformed to the subject’s functional space) was performed.

fMRI data preprocessing consisted of: EPI distortion correction using FUGUE (FMRIB’s Utility for Geometrically Unwarping EPIs) based on the acquired fieldmap; motion reduction with respect to the middle volume (reference) using FMRIB’s Linear Image Registration Tool (FLIRT), yielding 6 rigid-body motion parameters; and high-pass temporal filtering with a cut-off frequency of 0.01 Hz to remove slow drift fluctuations. Moreover, we regressed out the extended 24 motion parameters, based on the Friston 24-parameter model (Friston et al., 1996), together with WM and CSF signals obtained by averaging the BOLD time series across the respective masks, and the motion outliers (MO) identified using the FSL Motion Outliers tool with the dvars option thresholded at the 75th percentile plus 1.5 times the interquartile range (Power et al., 2012). Finally, spatial smoothing was performed using SUSAN tool (FSL’s SUSAN, S.M. Smith and J.M. Brady, 1997), employing a Gaussian kernel with a full width half maximum of 3.3 mm.

### 2.4 MRI data analysis

The framework used for data analysis is inspired by the work of Amico & Goñi (2018), in which principal component analysis (PCA) is applied in the connectivity domain and the FC matrices are then reconstrcuted using the number of principal components (PCs) that maximizes identifiability, i.e. how similar a subject is to him/herself and how different he/she is from other subjects. We extend this to a novel multilevel clinical fingerprinting framework in which identifiability is maximized over three levels: within-subject, within-session and group. The resulting FC fingerprints were then used for two separate analysis: computing the correlation among them and applying Network-Based Statistic (NBS) for within/between-group comparisons. Finally, the association between the results of both analysis and clinical features for the M group were investigated. The schematic pipeline representing the workflow employed to perform these steps is depicted in Figure 2. The code was implemented in MATLAB software version R2016b and is available in https://github.com/isesteves/multilevel-fingerprinting_migraine.

**Figure 2.**
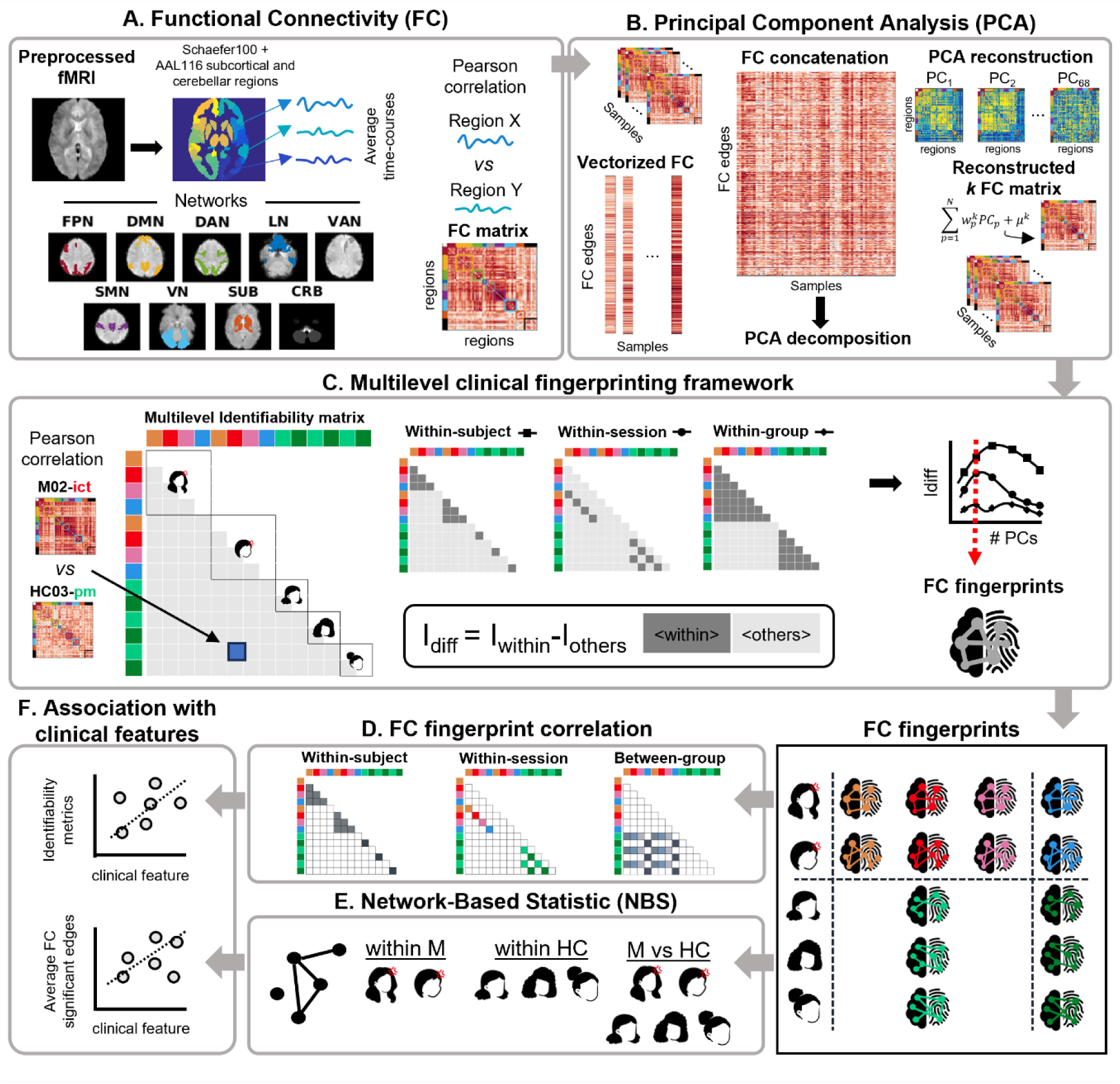
Pipeline scheme: A. Parcellation into regions of networks, based on the Schaefer atlas (100 regions, representing 7 networks) and subcortical and cerebellar regions from the AAL116 atlas: fronto-parietal (FPN), default mode (DMN), dorsal attention (DAN), limbic (LN), ventral attention (VAN), somatomotor (SMN), visual (VN), subcortical (SUB) and cerebellar (CRB). The parcellated data is used for FC computation, by calculating the Pearson correlation between the average time-courses of each pair of regions; B. The lower triangular matrix (excluding the diagonal) of each FC matrix was vectorized and they were all concatenated for PCA decomposition and reconstruction varying the number of PCs N, for each sample k; C. Multilevel clinical fingerprinting framework with PCs selection using a multilevel identifiability matrix corresponding to the Pearson correlation between vectorized FC matrices for all possible combinations of subjects/sessions (samples). In this matrix, for each level, I_within_ corresponds to the average of the elements in dark gray, I_others_ corresponds to the average of the elements in light gray and I_diff_ (%) corresponds to 100×(I_within_-I_others_). The FC fingerprints are obtained by reconstructing the data using only the selected number of PCs; D. FC fingerprint correlation was computed within-subject, within-session and between-group by averaging the correlation values per subject for each of these cases; E. Computation of Network-Based Statistic (NBS) for within- and between-group FC analysis. F. Association of identifiability metrics and average FC significant edges (found with NBS) with clinical features, using Spearman correlation.

#### 2.4.1 Functional connectomes (FC)

fMRI data was parcellated by computing region average time courses using Schaefer atlas (Schaefer et al., 2018) with 100 parcels, directly corresponding to the 7 intrinsic networks defined by (Thomas Yeo et al., 2011): fronto-parietal network (FPN), default mode network (DMN), dorsal attention network (DAN), limbic network (LN), ventral attention network (VAN), somatomotor network (SMN) and visual network (VN). Furthermore, given the importance of subcortical regions for migraine, as well as the cerebellum, two additional networks were retrieved from the AAL116 atlas (Tzourio-Mazoyer et al., 2002): one comprising the hippocampus, amygdala, caudate, putamen, pallidum and thalamus (SUB) and another one comprising 9 bilateral cerebellar regions and the vermis (CRB). Overall, the 9 networks encompassed a total of 130 regions (Figure 2.A). Care was taken to ensure that regions overlapping less than 50% with the fMRI data coverage were excluded for further analysis. This procedure led to the bilateral exclusion of 4 cerebellar regions: Crus II, Cerebellum7b, Cerebellum8 and Cerebellum9, for all subjects. The time series of each region were demeaned and bandpass-filtered from 0.01 to 0.1 Hz using a 2^nd^ order Butterworth filter. Symmetric FC matrices (130 regions×130 regions, 16900 edges) were generated by computing the Pearson correlation coefficient for each pair of regions.

#### 2.4.2 Principal Component Analysis

The lower triangular FC matrix (excluding the diagonal) was obtained for each subject and session, and vectorized (1×8385 edges). The individual vectors (10 M subjects × 4 M sessions = 40; 14 HC subjects × 2 HC sessions = 28; total = 68 samples) were concatenated across of all subjects and sessions into a single matrix (8385 edges×68 samples) (Figure 2.B). Subsequently, PCA was applied to obtain a number of PCs equal to the number of FC matrices in the dataset (i.e., 68 PCs). The PCs were ordered by decreasing variance explained, and the FC matrices was reconstructed by using an increasingly larger number of PCs, from 1 to 68. We expect that higher variance PCs will contain global FC information (session- and group-level), intermediate variance PCs will contain subject-level information, and the lowest variance PCs will capture noise/artifacts.

#### 2.4.3 Multilevel clinical fingerprinting

##### Multilevel identifiability matrix

Previous fingerprinting studies assessed subjects in at most two sessions (test and retest), either by evaluating them on two different days or by splitting single day data into two halves. In contrast, the present study assesses patients (M) in 4 timepoints and controls (HC) in 2 timepoints. To account for the larger number of sessions, we built a modified identifiability matrix by computing the Pearson correlation between all possible pairs of subjects/sessions (see Figure 2.C). This multilevel identifiability matrix is organized by subject: the first 40 rows/columns correspond to M, with each group of 4 being M-pre, M-ict, M-post and M-inter of each patient; the last 28 rows/columns correspond to HC, with each group of 2 being HC-pm and HC-postov. Hence, unlike the originally proposed identifiability matrix, this modified version is symmetric (68 samples x 68 samples). For ease of interpretation and to differentiate it from the original, we display our multilevel identifiability matrix by using only the corresponding lower triangular matrix.

##### Multilevel differential identifiability

The functional connectome fingerprinting framework is based on the premise (already substantiated by evidence) that a subject’s FC profile is more similar to their own FC profile assessed on a different day or task than to those of other subjects (Finn et al., 2015, 2017). Besides within-subject similarity, we extend the differential identifiability concept proposed by Amico & Goñi (2018) to consider also within-session and within-group similarity. The original framework introduced self-identifiability (I_self_) as the Pearson correlation between two sessions of the same subject (Amico & Goñi, 2018). To accommodate the different levels of similarity, we propose the use of within-identifiability (I_within_), a broader concept which corresponds to the average Pearson correlation between FC matrices belonging to the same subject, the same session, or the same group. Correspondingly, for each of the levels, the elements that are not included in I_self_ are considered I_others_, *i.e.*, average Pearson correlation between different subjects, different sessions, or different groups. The differential identifiability for each level is defined as the difference between these two terms: I_diff_ = 100×(I_within_-I_others_) (Figure 2.C). The higher the I_diff_, the more pronounced the individual fingerprint along that specific level. In this case, instead of maximizing only the within-subject I_diff_, we aim for a compromise that simultaneously maximizes within-subject, within-session and within-group I_diff_. Therefore, we manually selected the number of PCs used for the PCA reconstruction that provided the best trade-off across all I_diff_ levels. Having selected the number of PCs, we consider that the final reconstructed individual FC matrices correspond to the individual FC fingerprints.

##### FC fingerprint correlation

Upon obtaining the individual FC fingerprints, our goal was to interpret the results in the context of the longitudinal assessment of patients relative to controls. We extracted individual Pearson correlation values from the multilevel identifiability matrix computed on individual FC fingerprints, in order to average them for different comparisons and examine their distribution (see Figure 2.D). First, within subject, by separately analyzing M subjects (averaging 6 values for each subject, corresponding to combinations of 4 sessions, two by two) and HC subjects. Second, within session, considering all the points that corresponded to either M-pre, M-ict, M-post or M-inter for the M group and HC-pm and HC-postov, for the HC group. For each subject and session, we extracted the correlation values with all the other subjects in the same session and averaged them. Third, between groups, considering all the correlation values between each M session and the corresponding HC session: M-pre vs HC-pm, M-ict vs HC-pm, M-post vs HC-pm and M-inter vs HC-postov. For each subject in the M group and session, we extracted and averaged the correlation values with all the other subjects in the corresponding session of the HC group.

#### 2.4.4 Network-Based Statistic (NBS)

Using the previously obtained individual FC fingerprints for each subject and session, we employed Network-Based Statistic (NBS) to test for differences in the FC strength (Figure 2.E). This is a cluster-level tool for human connectome statistical analysis that models FC as a graph and controls for family-wise error rate (FWER) when performing mass univariate testing, which is available as a MATLAB toolbox (Zalesky et al., 2010).

#### 2.4.5 Association with clinical features

We investigated the association of FC identifiability and FC strength with relevant clinical features (Migraine Onset, Disease Duration, Attack Frequency, Attack Duration and Attack Pain Intensity), using Spearman correlation (Figure 2.F). Regarding identifiability, we evaluated the association with: 1) within-subject average correlation, 2) between-group average correlation for each M session; 3) between-group using the average correlation across all sessions for each patient. For FC strength, we calculated the average FC across all edges that showed significant differences when comparing M and HC.

### 2.5 Statistics

Due to the limited sample size, non-parametric statistical tests were used. Specifically, for comparisons within the M or HC group, we applied the signed rank Wilcoxon test. For comparisons between the M and HC groups, conducted specifically for matching sessions (M-pre, M-ict, M-post compared to HC-pm; M-inter compared to HC-postov), we employed the Wilcoxon rank sum test. The significance level was set to p-value = 0.05 and multiple comparisons were corrected for the false-discovery rate with the Benjamini-Hochberg method, using the Multiple Testing toolbox (Martínez-Cagigal, 2021).

The NBS detects network components that consist of connections surpassing a certain threshold. Subsequently, permutation testing is carried out to establish a p-value adjusted for FWER for each network, taking into account its size. The initial threshold for the test statistic was established at a t-value of 4.0, and 5,000 permutations were executed to identify a network component with a p-value < 0.05 after correction for FWER.

## 3. Results

### 3.1 Functional connectomes

The FC matrices obtained for all subjects and sessions are presented in Figure 3. The distributions display a shift towards positive correlation values and, for certain instances, a distinct between- or within-network pattern is discernible, as expected. Furthermore, several subjects exhibit a distinctive and consistent pattern across sessions despite some inter-session variability. However, across subjects within the same session, a clear pattern is not evident.

**Figure 3.**
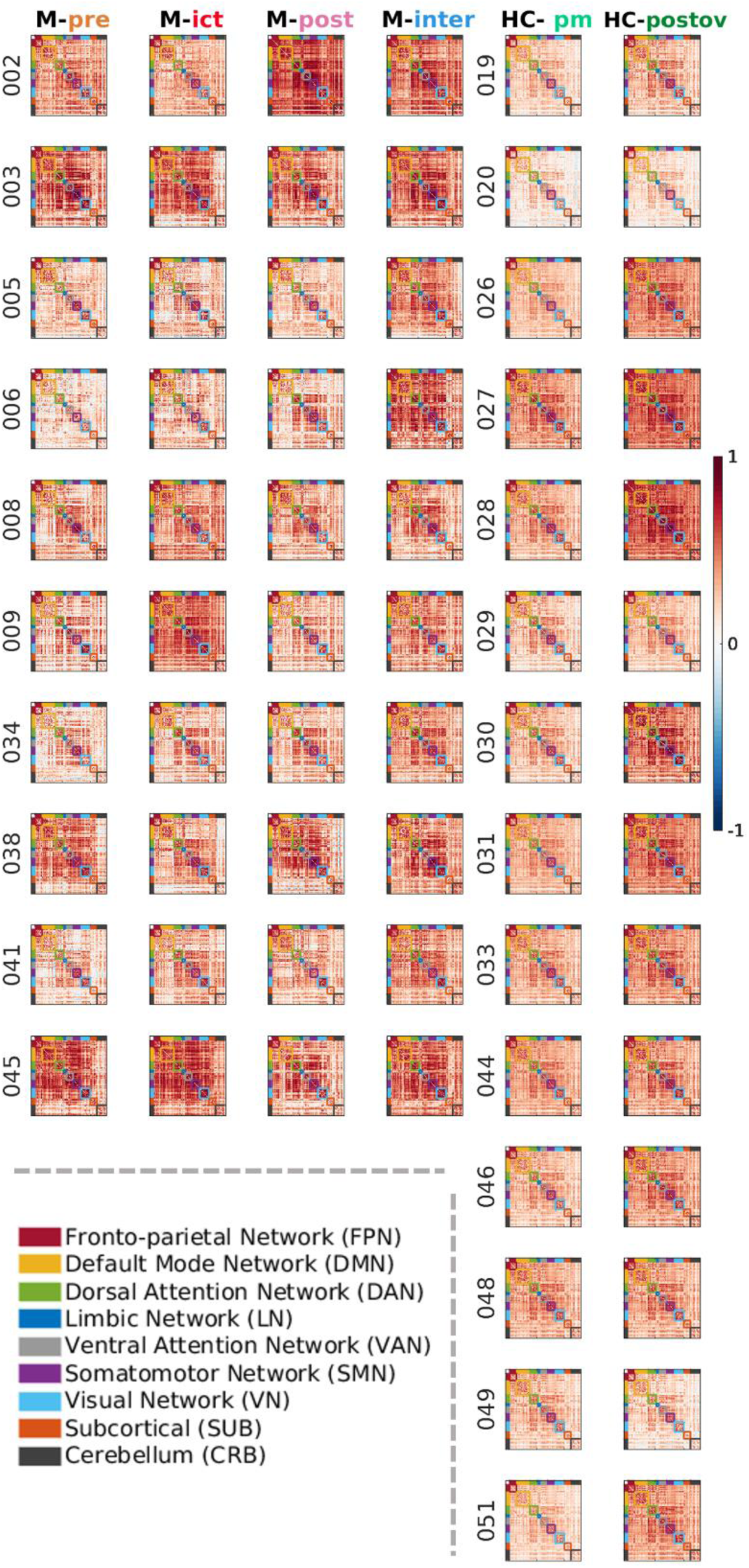
Functional connectivity matrices computed using Pearson correlation for each participant in the M group and HC group, in each session. In each FC matrix, the brain regions are ordered by network.

### 3.2 Multilevel clinical fingerprinting

#### Multilevel differential identifiability

The I_diff_ values computed for each level (within -subject, -session and -group) as a function of the number of PCs are presented in Figure 4.A, showing that the number of PCs that maximizes I_diff_ is not the same for all levels (within-subject: 24 PCs; within-session: 6 PCs; within-group: 36 PCs), as expected. Moreover, across all the numbers of PCs, I_diff_ values were larger within-subject than within-session and within-group. For our multilevel-informed selection of the number of PCs, since previous studies have focused on the subject, we used it as a starting point, focusing on its plateau region. In this region, we opted for a local maxima of the within-session and within-group levels, selecting a cutting-off point of 19 PCs. With this reconstruction, 80% of the variance of the original data was preserved. In Figure 4.B, we show the multilevel identifiability matrix after reconstruction with 68 PCs (100% of variance explained) and the selected 19 PCs (80% of variance explained), where specially the within-subject pattern is evidenced.

**Figure 4.**
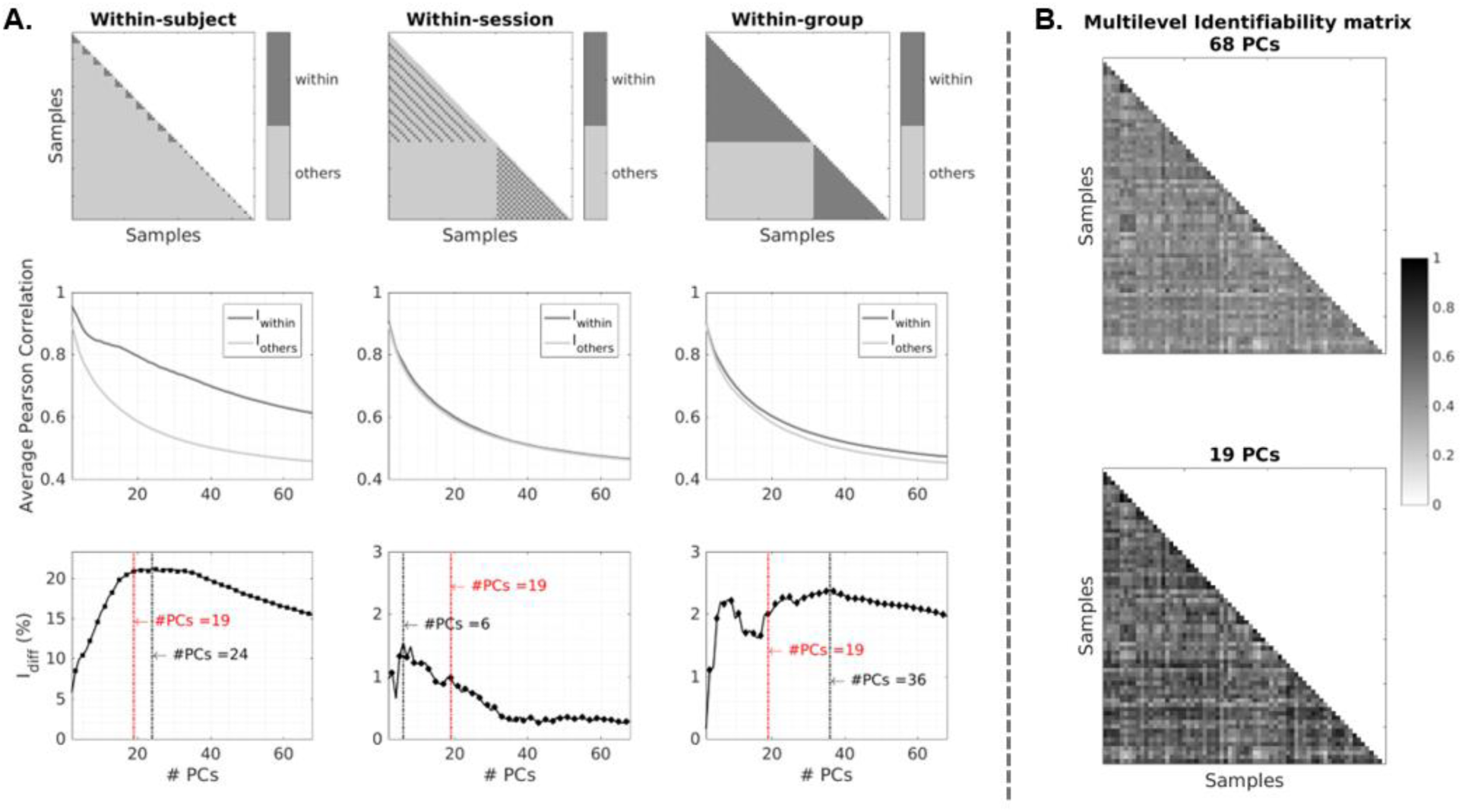
Multilevel Identifiability analysis: A. Multilevel templates for I_within_ (average correlation among equivalent samples) and I_others_ (average correlation among non-equivalent samples)(Top); I_within_ and I_others_ as a function of PCs (Center); I_diff_(%)=100x(I_within_-I_others_) as a function of PCs: 19 PCs (indicated in red) were chosen for reconstruction as a compromise between the values of all the levels (Bottom). B. Multilevel identifiability matrices computed using all PCs (68 PCs), and using only the selected ones (19 PCs), which shows localized increases in correlation values.

#### FC fingerprint correlation

The results of the FC fingerprint correlations are presented in Figure 5. At the within-subject level (Figure 5, left), no significant differences were observed between the M and HC groups. Although the M group exhibited a generally smaller dispersion, it is noteworthy that there were two outliers within this group, while the HC group did not show any. Within-session (Figure 5, center), significant differences were observed between HC-pm and M-pre. For the HC group, no significant differences were verified, whereas for the M group, M-pre was significantly different from M-post and M-inter, and M-ict was significantly different from M-inter. Thefore, during the M-pre, the FC of the M group not only deviates from that of the HC group, but also stands out as the most distinct among other migraine phases of the M group. In the between-group analysis (Figure 5, right), no significant differences were detected. Although there is a trend towards growing similarity between the HC and M groups throughout the migraine cycle, particularly peaking during the M-inter, the substantial variability prevents us from drawing conclusions.

**Figure 5.**
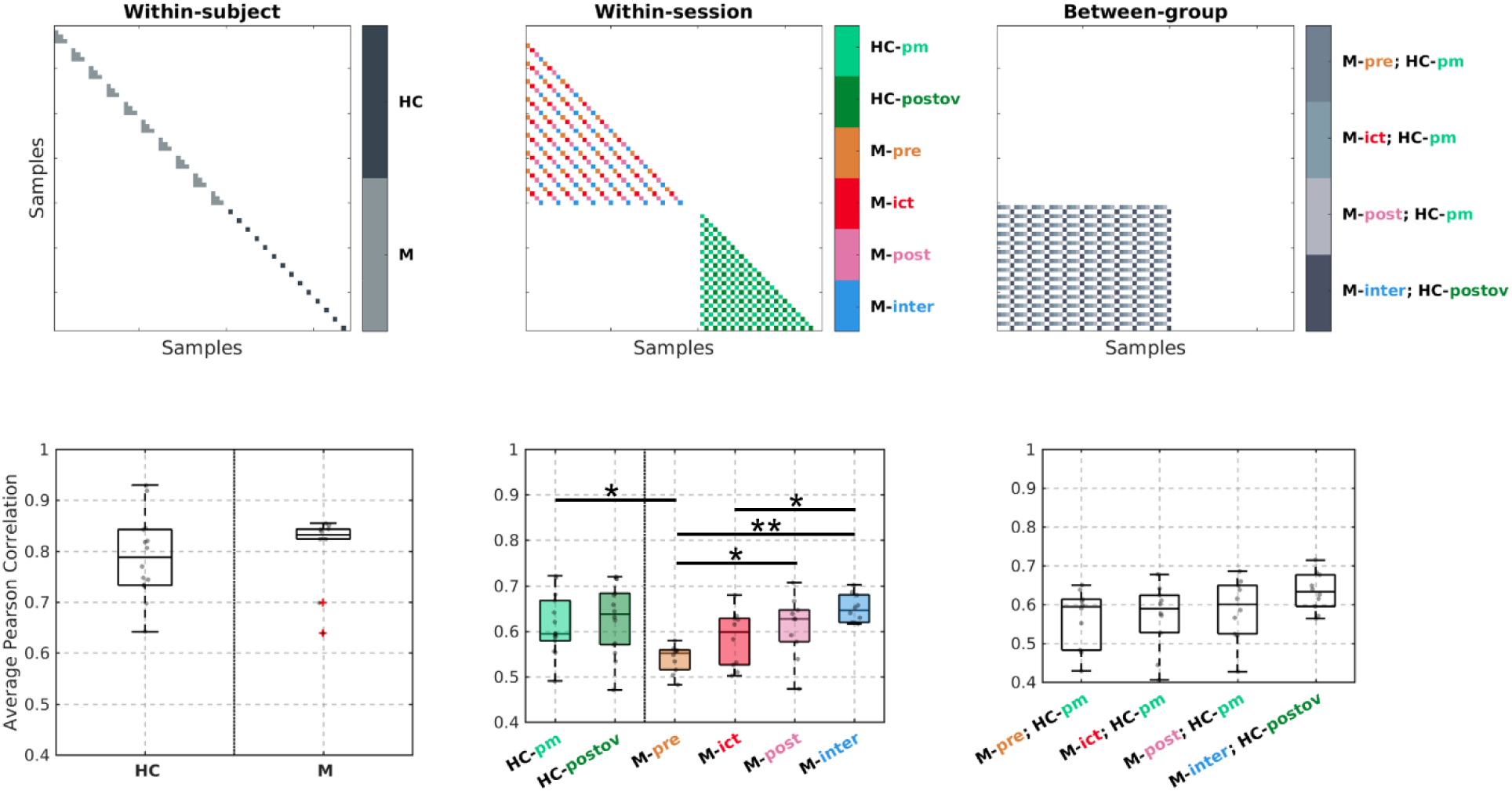
FC fingerprints average correlations: templates for computation (Top) and distributions across subjects (Bottom), for: within-subject (all pairwise session combinations for M; HC-pm vs HC-postov), within session (M/HC: all other M/HC during the same session), and between-group (only for M: all HC in the corresponding session) correlations. Significant differences are indicated; (Wilcoxon signed-rank test (1-sample) and Wilcoxon rank-sum tests (2-sample); Benjamini-Hochberg FDR correction; *p<0.05;**p<0.01).

### 3.3 Network-Based Statistic

The results for the analysis of FC strength within the M group (migraine phases within the same menstrual phase, i.e. preictal, ictal and postictal) and between-group (patients and controls in matching phases, i.e., preictal, ictal and postictal compared to perimenstrual; interictal compared to postovulation) are shown in Figure 6. Comparing the HC and M groups, in the corresponding phases, there were no significant differences between M-postov and HC-inter but M-ict and M-post both showed signicantly higher FC compared to HC-pm. While M-ict had higher FC primarily in connections involving the SMN, VN, VAN, DAN, and CRB, M-post had higher FC in fewer connections mostly the VN and attentional networks.

**Figure 6.**
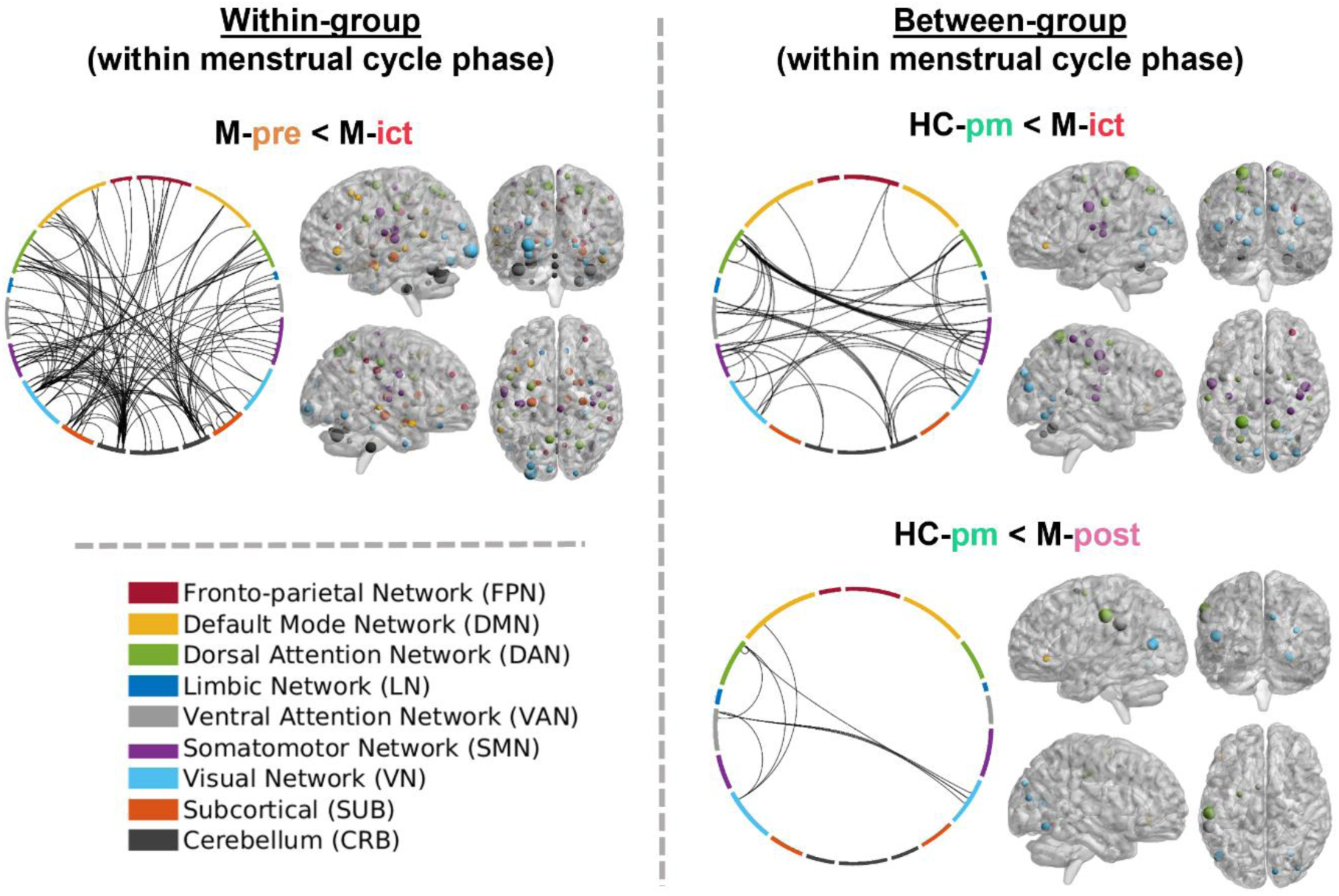
FC analysis within-group and within menstrual cycle phase (Left) and between-group (Right), using NBS (based on extent, cluster threshold = 4; α = 0.05; 5000 permutations). For each comparison, two representations are shown: a chord diagram with the edges that significantly differ (generated with NiChord Python toolbox (Bogdan et al., 2023)) and four brain views showing the nodes that contribute to significantly different edges (generated with BrainNet Viewer MATLAB toolbox (Xia et al., 2013)). For each comparison, the node size is scaled according to the node degree, which corresponds to the sum of the number of significantly different edges linked to that node. The networks are represented by colors, as indicated in the legend.

Regarding the within-group (patients) comparisons, for the same menstrual phase, M-ict showed higher FC compared to M-pre, mainly between the CRB and other networks (specially SMN, VAN, DAN, VN and SUB). It should be noted that, when considering 68 PCs instead of only the selected 19 PCs, the differences between M-pre and M-ict did not survive the cluster threshold (results not shown). This indicates that the PCA reconstruction step was advantageous for refining this analysis.

Additionally, the results obtained by comparing the different phases within each group across different menstrual phases (i.e., preictal, ictal and postictal compared to interictal for patients; perimenstrual compared to postovulation for controls) are presented in Figure S1 (Supplementary Material). For the controls, FC was lower in the perimenstrual relative to the postovulatory phase, mainly in connections within the VN and between the VN and other networks (DAN, DMN, CRB, and FPN). For the patients, FC was also lower in the peri-ictal phases (M-pre, M-ict, M-post) compared to the interictal phase, with connections between VN and DMN as well as DAN and DMN being common across the three phases. These findings highlights the signifiant impact of the menstrual cycle in the FC strength, and hence the importance of controlling for it when studying the migraine cycle. For this reason, within-patients comparisons were considered only witin the same mentrual cycle phase in the main Figure 6.

### 3.4 Association with clinical features

The average FC in edges significantly different between M-post and HC-pm was negatively correlated with the Attack Frequency (r_s_ = -0.83, p_FDRcorr_ < 0.05). On the contrary, it was positively correlated with the Migraine Onset, at the uncorrected significance level, but failed to surpass the multiple comparisons correction threshold (r_s_ = 0.66, p_uncorr_ < 0.05). There were no significant correlations of differential FC identifiability measures or FC strength with any of the other clinical features.

## 4 Discussion

We report the first case-control longitudinal study of brain FC across all phases of the migraine cycle, employing a novel multilevel clinical fingerprinting approach to retrieve individual FC fingerprints of patients and their healthy controls. We found decreased FC identifiability of patients in the preictal phase compared with controls, which increased after the attack reaching normal values in the interictal phase. Moreover, we found that FC fingerprints of patients exhibited increased strength in the ictal and postictal phases but were otherwise comparable to those of healthy controls in the corresponding menstrual phases.

### Multilevel clinical fingerprinting framework

The multilevel fingerprinting approach allowed to capture representative FC fingerprints, by excluding redundant/artifactual information. We observed increased FC heterogeneity in the preictal/ictal phases, which decreased with the progression of the attack. Furthermore, the preictal phase was the only one during which migraine patients were significantly more heterogeneous compared to healthy controls. When not experiencing symptoms, patients were more homogeneous, and this homogeneity did not differ from the one presented by controls. This is partially in line with what was observed in previous fingerprinting studies addressing disease, where there was also more patient heterogeneity for amnesic mild cognitive impairment (Sorrentino et al., 2021), amyotrophic lateral sclerosis (Romano et al., 2022), Parkinson’s disease (Troisi Lopez et al., 2023) and multiple sclerosis (Cipriano, Troisi Lopez, et al., 2023). Nevertheless, these studies also observed larger intra-subject variability for patients, which was not the case for migraine. It is noteworthy, though, that they all employed MEG instead of fMRI, and focused on a very different time scale (test and re-rest sessions were measured on the same day, with only 1 min break between them). Furthermore, in our case within-subject correlation for the M group was not significantly correlated with any of the clinical features.

Our between-group average correlation is similar to the I_clinical_ metric proposed by (Sorrentino et al., 2021) except for the fact that in their case and related work (Cipriano, Troisi Lopez, et al., 2023; Romano et al., 2022; Troisi Lopez et al., 2023), there were only two sessions and they were considered equivalent. In our case, it is relevant to assess each of them individually since we expected differences at each phase of the migraine cycle. Nevertheless, in our case it was not possible to establish an association between this metric and any of the clinical features, possibly because the other studies assessed a larger sample.

Concerning the menstrual cycle, there is currently one study (available as a preprint, not yet peer-reviewed) evaluating fingerprints across three phases: peri-ovulatory, mid-luteal and early follicular (Cipriano et al., 2023). They observed that the identifiability remained consistent across phases. Nonetheless, a more detailed edgewise analysis revealed that the connectomes during the peri-ovulatory and mid-luteal phases showed less stability. Although the peri-ovulatory phase and early follicular phase in that study partially coincide with the post-ovulation and perimenstrual phases defined in the present study, the methodological differences hinder us from establishing strong connections between the two studies.

### FC fingerprints analysis

#### FC changes across the menstrual cycle

Regarding FC investigated with NBS applied to FC fingerprints, there were differences between HC phases, including all networks specially connections with bilateral VN, and involving SUB regions to a lesser extent compared to other networks. The functional and structural changes occurring across the menstrual cycle have started to gain attention only recently. Despite the methodological challenges as well as the variety of scanning protocols and timings that have been used, the comprehensive review performed by (Dubol et al., 2021) shows FC changes across the cycle in regions such as the hippocampus, amygdala, anterior cingulate cortex, insula, inferior parietal lobule, and prefrontal cortex. Nevertheless, it did not include studies comparing the early luteal phase (post-ovulation) and late luteal phase or early follicular phase (perimenstrual), probably because the levels of ovarian hormones (progesterone and estradiol) are not expected to differ a lot during these periods. In our case, although it was not possible to measure hormone levels, considering the timings of post-ovulation and perimenstrual sessions, it is more likely that hormonal differences concern higher estradiol and progesterone levels in the post-ovulation phase, compared to perimenstrual. In general, higher levels of these hormones were mostly linked with increased FC (Dubol et al., 2021; Pritschet et al., 2020), which is in line with our results, considering that we found significantly increased FC in HC-postov compared to HC-pm, and we did not find any evidence of decreased FC.

#### FC changes across the migraine cycle (across menstrual cycle phases)

Considering the differences across the menstrual cycle for HC, differences between the peri-ictal phases and the interictal phase should be carefully interpreted. Compared to M-inter, all peri-ictal phases exhibited decreased FC concerning several networks. M-pre shows decreased FC mostly bilaterally for VN, SMN, and DAN, the right FPN and DMN, and the vermis (CRB). Another longitudinal experiment had found increased FC in the preictal phase compared to the interictal phase between the nucleus accumbens and the amygdala, hippocampus and parahippocampal gyrus (associated with limbic function) (Schulte, Menz, et al., 2020). The nucleus accumbens and parahippocampal gyris were not included in this analysis, though we verified significant differences between subcortical regions, yet mostly with the FPN. M-ict shows decreased FC mostly involving connections between the DMN, DAN, VN and right SMN. Amin et al. (2018) found evidence of decreased FC between the right thalamus and structures from the ipsilateral SMN, though there was also increased FC between the right thalamus and contralateral structures from the LN, DAN, SMN and VAN. Moreover, the FC decrease between the SMN and VAN (posterior insula) observed by (Araújo et al., 2023) was not replicated, though other connections involving the SMN or VAN showed decreased FC in our case. Decreased FC within limbic structures and with the hypothalamus has also been observed (Stankewitz et al., 2021; Stankewitz & Schulz, 2022). Furthermore, M-post shows decreased FC bilaterally for connections involving SMN and VN, as well as left DAN and DMN, yet no differences were found in the existing literature. Considering the decreased FC during HC-pm compared to HC-postov, we would expect the FC of more connections to be decreased when comparing the peri-ictal phases with M-inter.

#### FC changes across the migraine cycle (within menstrual cycle phase)

Across the migraine cycle, comparing the peri-ictal sessions, there were significant differences between M-pre and M-ict, though not when comparing it to M-post. Compared to M-ict, M-pre shows decreased FC for connections among all networks, bilaterally. Furthermore, cerebellar regions and part of the left VN (particularly comprising the occipital pole) are greatly involved in these connections. For CRB, most connections are to the SMN (especially left), although the left SUB, VN, VAN and DAN are highly involved as well. Stankewitz & Schulz (2022) also found decreased VN and CRB FC, using a linear mixed-effects model to explain the trajectory (over several migraine cycle points) of the FC of regions belonging to 48 networks defined by group Independent Component Analysis. However, contrary to our results, they also observed increased FC in 22 networks, mainly involving regions belonging to VN, CRB, SMN, VAN, DMN and LN. Nevertheless, contradictory results could be attributed to methodological differences and the fact that they studied a mixed sample with and without aura, whereas our sample is restricted to migraine without aura (usually occurring during the preictal phase).

#### FC differences between M and HC (within menstrual cycle phase)

Comparing M with HC during the same menstrual cycle phase, there were no differences between M-inter and HC-postov. Regarding the interictal phase, the literature shows conflicting results. A systematic review conducted by (Skorobogatykh et al., 2019) pointed out that, although FC studies had found differences in more than 20 networks, these were not reproducible and there was no common migraine pattern. The more recent review by (Schramm et al., 2023), investigating FC among other methods, focused on the insula, brainstem, thalamus, hypothalamus, limbic areas and several functional networks. Despite the fact that several studies report differences, there are also several inconsistencies, such as opposing results and lack of reproducibility. Moreover, a recent study examined a large sample suffering from migraine with aura and did not find any resting state differences between patients and controls and the meta-analysis performed by (Chen et al., 2022) did not find consistent FC differences. Nevertheless, another recent meta-analysis focused only on the DMN found significant interictal differences compared to healthy controls (Hu et al., 2023). Considering the prevalence of menstrual-related migraine and the fact that most studies have considerably larger percentage of female participants, some inconsistencies may be due to inadequate control of hormonal factors.

Nevertheless, comparing to HC-pm, both M-ict and M-post show differences, while M-pre does not. For M-ict, FC is increased mostly between left DAN and right SMN, left VAN and right CRB, and also other connections involving the VN, DAN, VAN and SMN, bilaterally. Regions belonging to the DAN (especially the parietal lobule), to the SMN (comprising the precentral/postcentral gyrus), and the CRB correspond to nodes with more significantly increased connections. Other studies had found differences between the ictal phase and controls: while there was a decrease in FC between FPN and both DAN and VAN (Coppola et al., 2016), there was also an increase of FC within the DMN and between the DMN and VAN (Coppola et al., 2018). For M-post, FC was increased mostly between left attentional networks (VAN and DAN) and VN bilaterally, between the left attentional networks themselves, and the right DMN and ipsilateral VN. In this case, the regions with the highest number of connections belong to the DAN and VAN, both overlapping with the supramarginal gyrus, and the VN. Furthermore, when comparing HC-pm to both M-ict and M-post, although with different connections, there are several regions sharing an increase in FC, mainly in the left DMN, DAN and VAN, as well as bilateral VN. Concerning the preictal phase, significant decreases in FC compared to healthy controls had been previously observed during painful stimulation (Marciszewski et al., 2018) and resting state (Meylakh et al., 2018) for the brainstem, which we did not include. This structure was not investigated in our case since our acquisition and analysis were not optimized in that sense, as it would be ideal due to its location.

#### Summary

Thus, we hypothesize that the occurrence of migraine may lead to an abnormal increase in FC during peri-ictal phases after the migraine attack has initiated (following the preictal phase), when compared to the FC expected for a normal perimenstrual phase. Therefore, experiencing a migraine attack may lead to a quicker reset of FC perimenstrually that would not occur as fast considering only normal hormonal fluctuations across the menstrual cycle. Nevertheless, to confirm it, further studies should be performed with a longitudinal assessment of migraine patients during menstrual cycles with and without the occurrence of migraine, ideally accompanied by hormonal levels measurement.

### Limitations

Despite recognizing the value and uniqueness of this dataset, we must acknowledge several limitations in our study. The sample size remains limited. Additionally, it would have been ideal for migraine patients to abstain from using hormonal contraception, as this introduces an additional variable. Even for women with natural menstrual cycles, incorporating hormonal level measurements could enhance the interpretation of FC fluctuations throughout migraine/menstrual cycles, potentially isolating effects directly associated with migraines.

Moreover, there exists intra-subject variability in migraine attacks, and the data was not consistently recorded from the same migraine cycle, leading to potential differences in hormonal factors and pain attack intensity. We did not register the timing of interictal sessions in relation to the last/next ictal session, which could have provided valuable insights.

While several studies have identified changes in the brainstem, we excluded it from our analysis due to the need for a specific acquisition setting and susceptibility to physiological noise. Further research is imperative to explore additional factors in-depth.

### Novelty and impact

Limited research has delved into the FC across the migraine cycle, as most studies focus on the interictal phase only. Challenges in studying phases around attacks, attributed to logistical hurdles and patient discomfort, have hindered thorough investigation, despite their crucial role in understanding the complete migraine cycle. Longitudinal assessments are scarce, with some focusing only on the ictal and interictal sessions, or different scans of the same cycle phase. Notably, some studies collected longitudinal data for all cycle phases, but their controls were lacking. Furthermore, most studies include mostly or only female patients, though most of them do not comment on the menstrual cycle phase or, if they do, they just generally mention not having collected data during the menstrual period. In summary, the existing functional connectivity studies exhibit a fragmented landscape, with few studies covering the full migraine cycle longitudinally and those that do it lacking comprehensive control comparisons. Therefore, we believe this study brings a valuable contribution by addressing the gaps in existing research, offering a more comprehensive and nuanced exploration of functional connectivity across the entire migraine cycle with attention to subject variability, control comparisons and the menstrual cycle of female participants.

## 5 Conclusions

We determined the influence of the migraine cycle on individual functional connectome fingerprints in a longitudinal study for patients with episodic menstrual or menstrual-related migraine without aura across the four phases of the migraine cycle and their healthy controls in corresponding phases of the menstrual cycle. Our novel multilevel clinical connectome fingerprinting approach enabled the detection of increased variability among patients, beyond menstrual effects, especially around migraine attacks when patients also differ more from controls. These FC fingerprint variations across the migraine cycle could potentially pave the way for tailoring treatment strategies based on individual patterns. To our knowledge, this work represents the first case-control longitudinal fMRI study across the whole migraine cycle, tackling a major cause of disability worldwide by contributing to developing connectome-based migraine biomarkers.

## Acknowledgements

This work was supported by LARSyS funding (DOI: 10.54499/LA/P/0083/2020, 10.54499/UIDP/50009/2020, and 10.54499/UIDB/50009/2020), grant PRR -02/C05-i01.02/2022 (BL307/2023_IST-ID) and FCT grants PD/BD/150356/2019, PTDC/EMD-EMD/29675/2017, LISBOA-01-0145-FEDER-029675.

## Notes

### Competing Interest Statement

The authors have declared no competing interest.

