## Supplementary Figure 1 for "Multilevel clinical fingerprinting: uncovering longitudinal changes in the functional connectome of the brain along the migraine cycle"

### Supplementary material

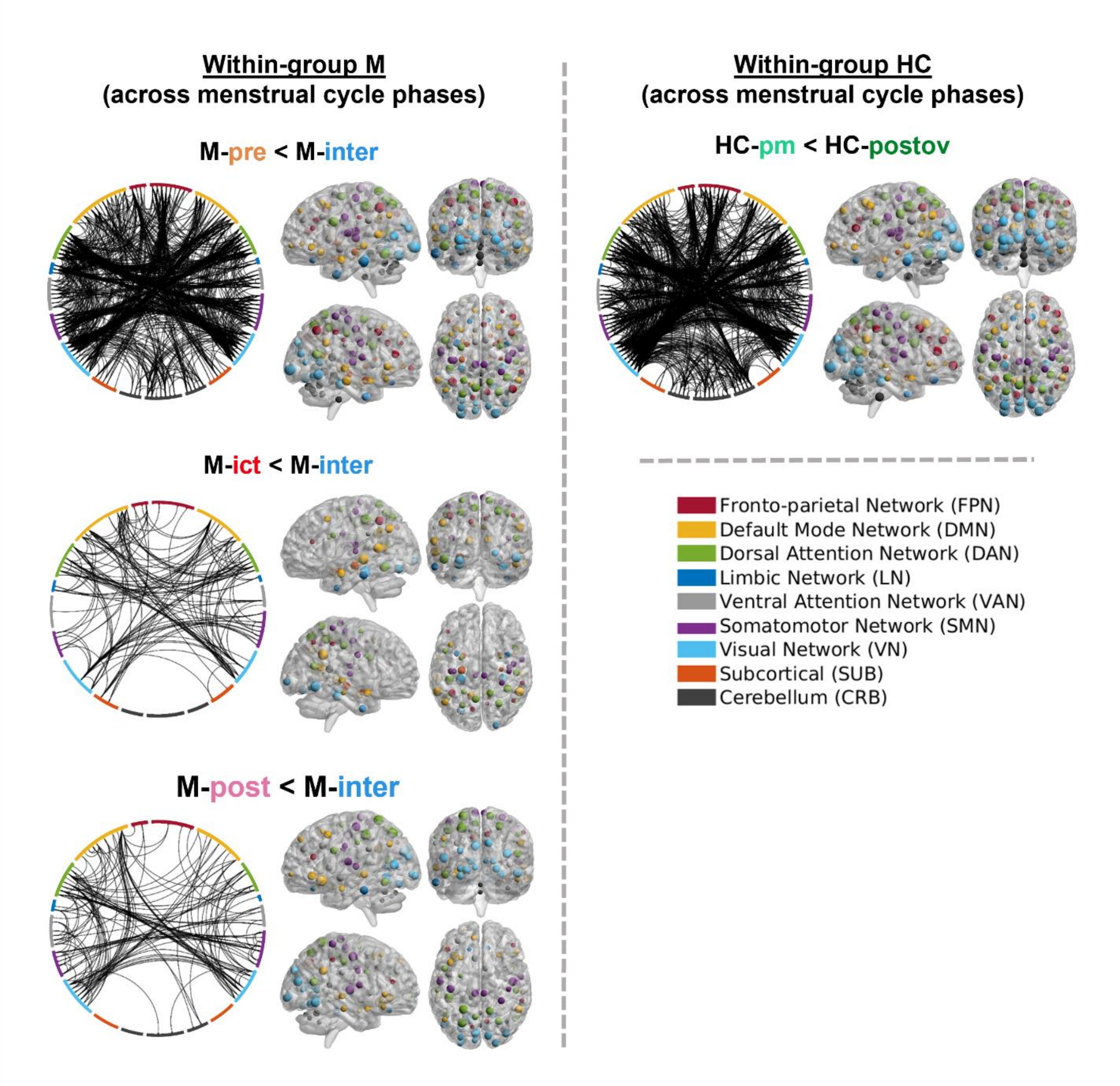


*Figure S1 - FC analysis within-group across menstrual cycle phases for the M group (Left) and the HC group (Right), using NBS (based on extent, cluster threshold = 4; α = 0.05; 5000 permutations). For each comparison, we present: a chord diagram with the edges that significantly differ (generated with NiChord Python toolbox (Bogdan et al., 2023)) and four brain views showing the nodes that contribute to significantly different edges (generated with BrainNet Viewer MATLAB toolbox (Xia et al., 2013)). For each comparison, the node size is scaled according to the node degree, which corresponds to the sum of the number of significantly different edges linked to that node. The networks are represented by colors, as indicated in the legend.*
